# PathoBench: an open community-driven benchmark registry for pathogen bioinformatics tools

**DOI:** 10.64898/2026.07.16.739016

**Authors:** Yibo Dong, Na Li, Calin B. Chiribau, Molly Mitchell, Xiaoming Liu, Andy D. Perkins

**Affiliations:** Bureau of Public Health Laboratories, Florida Department of Health, Jacksonville, FL, USA; Mayo Clinic, Jacksonville, FL, USA; Center for Global Health and Infectious Diseases Research and USF Genomics Program, College of Public Health, University of South Florida, Tampa, FL, USA; Department of Computer Science and Engineering, Mississippi State University, Starkville, MS, USA

## Abstract

**Summary:** The proliferation of pathogen bioinformatics pipelines has outpaced the community’s ability to compare them on common ground. Self-reported performance numbers, ad-hoc evaluation datasets, and inconsistent metrics make pipeline selection difficult for clinical and public-health researchers. We present **PathoBench**, an open web platform that addresses this gap through three coordinated mechanisms: (i) a curated registry of 26 standard benchmark datasets across 10 human pathogens, each with persistent identifiers and direct download links; (ii) pathogen-specific evaluation metrics that submissions must report, allowing direct head-to-head comparison only on the same dataset; and (iii) a credibility framework combining mandatory dataset attestation, ORCID-linked attribution, public peer comments, and administrator verification. As a case study, four published *Mycobacterium tuberculosis* drug-resistance pipelines were evaluated against the WHO TB mutation catalogue, demonstrating the framework’s discriminating power. PathoBench is open for community contributions across all ten supported pathogens.

**Availability and implementation:** PathoBench is freely available at https://pathobench.vercel.app. Source code is released under the MIT license at https://github.com/BPHL-Molecular/pathobench. The platform requires no installation for end users; programmatic access is available via a Supabase REST API.

**Supplementary information:** Supplementary data are available at *Bioinformatics* online.

## 1 Introduction

Next-generation sequencing has transformed pathogen surveillance, diagnostics, and outbreak response. To analyze the resulting genomic data, more than 30,000 bioinformatics tools have been released (Gauthier et al., 2019), with hundreds focused on individual pathogens of public-health importance. Faced with this abundance, researchers struggle to select tools that are accurate, well-maintained, and appropriate for their use case.

The conventional remedy — independent comparison studies (Allali et al., 2017; Dong et al., 2021) — provides useful snapshots but cannot keep pace with new releases, and inter-study comparisons are confounded by inconsistent test data and metrics. Standardized benchmarking initiatives have emerged in adjacent fields (Teng et al., 2016; Meyer et al., 2021) but no equivalent exists for the heterogeneous landscape of human pathogen bioinformatics, where each disease imposes distinct requirements: drug-resistance mutations differ between *M. tuberculosis* and HIV; lineage taxonomy is unique to SARS-CoV-2; and APOBEC3-driven mutation patterns are specific to recent MPXV evolution.

We present PathoBench, a web platform that converts pipeline benchmarking from a one-off effort into a continuously growing community resource. It contributes three design elements that, together, distinguish it from prior efforts: a curated registry of standard reference datasets, pathogen-specific yet uniformly structured metrics, and an explicit credibility framework that handles open submissions without sacrificing data quality.

## 2 Implementation

### 2.1 Architecture

PathoBench follows a serverless three-tier architecture (Figure 1a). The frontend is a React 18 single-page application built with Vite and Tailwind CSS; the backend uses Supabase, providing PostgreSQL storage with row-level security policies, magic-link authentication, and a REST API. Anti-abuse protection is provided by hCaptcha. Static assets are served via Vercel; the platform requires no maintained server infrastructure and is free to operate at the academic scale (estimated capacity ≥250,000 submissions on free tiers).

**Figure 1.**
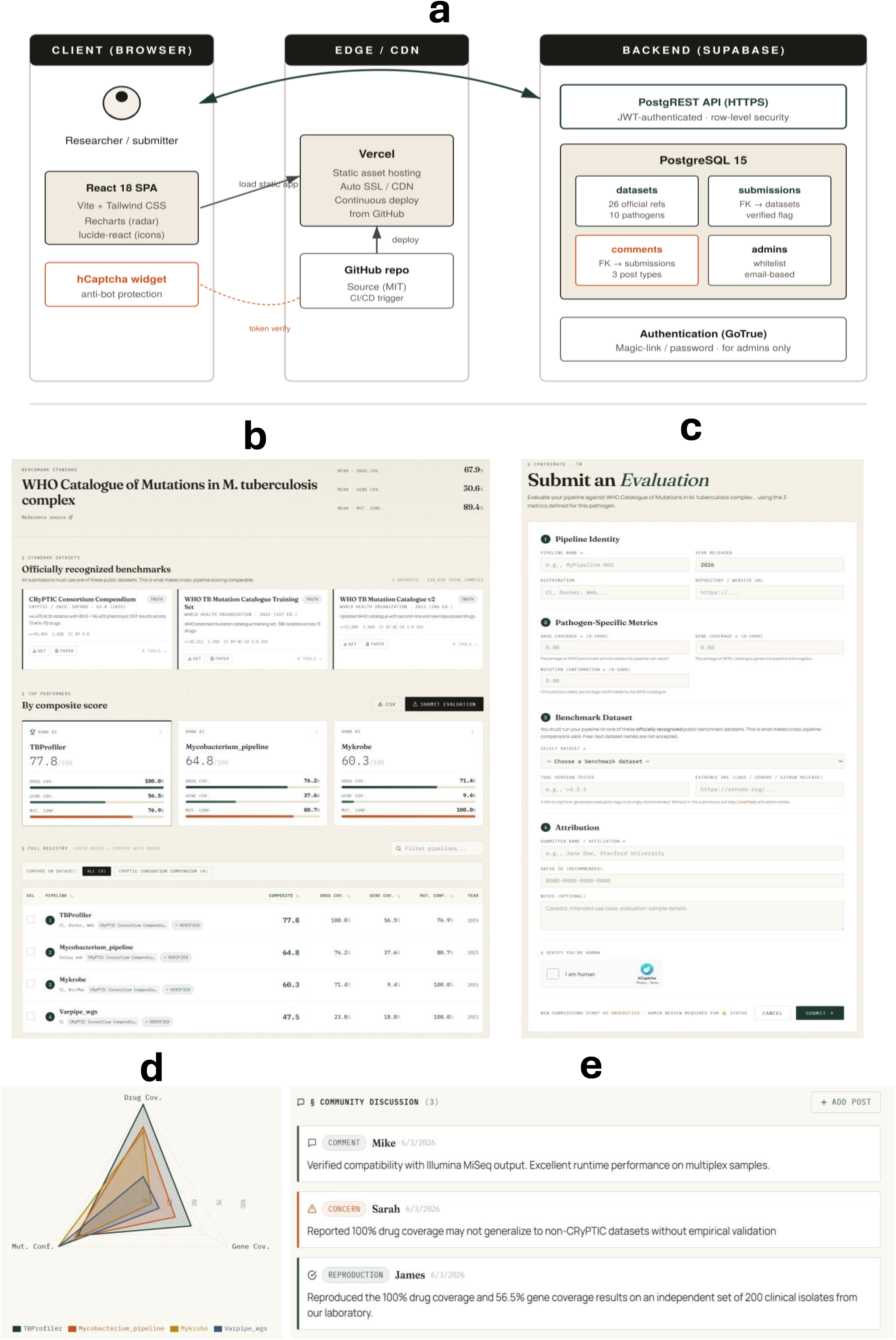
PathoBench platform interface and case study. (a) Architecture: a React frontend communicates with a Supabase backend (PostgreSQL + Auth + REST API) over HTTPS; static hosting is provided by Vercel; submissions are protected by hCaptcha. (b) Pathogen landing page for *M. tuberculosis*, showing the three registered benchmark datasets and the full pipeline leaderboard. (c) Submission workflow enforcing mandatory dataset selection from the registered list. (d) Comparative radar plot of the four TB drug-resistance pipelines on the WHO 2nd-edition catalogue, generated automatically from the leaderboard. (e) Public discussion thread on a submission, supporting comment, concern, and reproduction post types.

### 2.2 Standard datasets as first-class entities

A central design choice is that every submitted evaluation must reference a benchmark dataset registered in the platform. The initial release seeds 26 datasets curated from authoritative sources across ten pathogens (Table 1), each annotated with curator, sample count, license, DOI, and download URL. Examples include the CRyPTIC Consortium compendium for *M. tuberculosis* (Walker et al., 2022; n=44,405 isolates), the Pango designation set for SARS-CoV-2 (Rambaut et al., 2020), and MalariaGEN Pf7 for *Plasmodium falciparum* (MalariaGEN et al., 2023; n=20,864 samples). Datasets can be added or updated by platform administrators through a web interface; database-level foreign-key constraints prevent submissions referring to non-existent datasets.

**Table 1.** Standard benchmark datasets registered at platform launch (n = 26)

| Pathogen | Dataset | Curator | Samples | Reference |
| --- | --- | --- | --- | --- |
| *M. tuberculosis* | CRyPTIC v2 | CRyPTIC Consortium | 44,405 | Walker et al. 2022 |
| *M. tuberculosis* | WHO Catalogue v1 | WHO | 38,215 | WHO 2021 |
| *M. tuberculosis* | WHO Catalogue v2 | WHO | 52,000 | WHO 2023 |
| HIV-1 | Stanford HIVdb PR/RT | Stanford U. | 5,894 | Rhee et al. 2017 |
| HIV-1 | Stanford HIVdb Integrase | Stanford U. | 2,380 | Tzou et al. 2020 |
| HIV-1 | NIH PEPFAR EQA | NIH | 200 | – |
| HBV | HBVdb | INSERM Lyon | 120,000 | Hayer et al. 2013 |
| HBV | Stanford HBVdb | Stanford U. | 1,850 | Tan et al. 2007 |
| HCV | LANL HCV | LANL | 350,000 | – |
| HCV | ICTV references | ICTV | 8 | Smith et al. 2019 |
| HCV | HCV-GLUE test panel | MRC-Glasgow CVR | 1,245 | Singer et al. 2018 |
| HAV | HAVNET reference | RIVM | 425 | Enkirch et al. 2020 |
| HAV | NCBI HAV RefSeq | NCBI | 1,820 | – |
| SARS-CoV-2 | Pango designation | cov-lineages | 18,500 | Rambaut et al. 2020 |
| SARS-CoV-2 | Nextclade reference | Nextstrain | 850 | Aksamentov et al. 2021 |
| SARS-CoV-2 | GISAID curated | GISAID | 12,000 | – |
| MPXV | Pathoplexus MPXV | Pathoplexus | 5,200 | – |
| MPXV | Nextclade hMPXV | Nextstrain | 420 | – |
| Influenza | NCBI IVR | NCBI | 850,000 | Bao et al. 2008 |
| Influenza | GISAID EpiFlu | GISAID | 35,000 | – |
| Influenza | WHO CC panel | WHO | 1,500 | – |
| Ebola | Nextstrain Ebola | Nextstrain | 1,581 | Hadfield et al. 2018 |
| Ebola | NCBI Ebolavirus | NCBI | 195 | – |
| *P. falciparum* | MalariaGEN Pf7 | MalariaGEN | 20,864 | MalariaGEN et al. 2023 |
| *P. vivax* | MalariaGEN Pv4 | MalariaGEN | 1,895 | MalariaGEN et al. 2022 |
| *Plasmodium spp.* | PlasmoDB | VEuPathDB | 12 (genomes) | Aurrecochea et al. 2009 |

Datasets are dynamically loaded from the platform database; the current registry can be exported at any time via the Admin → Datasets CSV button.

This constraint is what enables valid head-to-head comparison: leaderboards can be filtered to show only pipelines evaluated on the same dataset, eliminating the most common confounder in informal benchmarks.

### 2.3 Pathogen-specific metrics

For each of the ten supported pathogens, PathoBench defines three to five evaluation dimensions that reflect that pathogen’s analytical priorities (Supplementary Table S1). For TB, these are drug coverage, gene coverage, and mutation confirmation against the WHO catalogue (WHO, 2023); for HIV, DRM concordance, drug-class coverage, minor-variant sensitivity, and score concordance against Stanford HIVdb. Submissions report all metrics as percentages, and a composite score is computed as the mean of pathogen-specific metrics. Cross-pathogen comparison is intentionally not supported.

### 2.4 Credibility framework

Open submission introduces an obvious tension between accessibility and reliability. PathoBench addresses this through four mechanisms operating in concert:

1. **Mandatory dataset attestation**. Free-text dataset descriptions are rejected; submitters must select from the registered list, providing a fixed denominator for comparison.
2. **Attribution**. Submissions request an ORCID identifier and link to an evidence URL (e.g. a Zenodo deposit or GitHub release containing machine-generated evaluation logs). Both fields render in the public record.
3. **Tiered verification**. Every submission is created with verified status set to false (displayed as a red “Unverified” badge). Designated administrators can mark submissions verified after reviewing evidence; verified entries display prominently and sort first on leaderboards.
4. **Public discussion**. Each submission carries a community discussion thread supporting three post types — comment, concern, and reproduction — enabling peer scrutiny analogous to post-publication review. Administrators can hide abusive content but cannot delete legitimate concerns.

These mechanisms make the platform’s reliability emerge from transparency rather than from gatekeeping.

## 3 Demonstration: TB drug-resistance pipelines

To illustrate PathoBench, we evaluated four published *M. tuberculosis* drug-resistance prediction pipelines — TBProfiler (Phelan et al., 2019), Mykrobe (Hunt et al., 2019), Varpipe_wgs (CDC), and Mycobacterium_pipeline (Sciensano) — against the 2nd-edition WHO TB mutation catalogue (WHO, 2023). Three metrics are computed: **drug coverage** (the proportion of WHO-listed antimicrobials reported by the pipeline), **gene coverage** (the proportion of resistance genes in the WHO catalogue covered by the pipeline), and **mutation confirmation** (the proportion of pipeline-called mutations that can be confirmed in the WHO catalogue). Results are reproduced in the live registry at https://pathobench.vercel.app/#tb (Figure 1d).

The four pipelines show clearly differentiated profiles. TBProfiler reports the broadest antimicrobial coverage (100%) and highest gene coverage (56.5%), making it well-suited to surveillance contexts where breadth is essential. Varpipe_wgs reports the narrowest antimicrobial coverage (23.8%) but achieves 100% mutation confirmation against the WHO catalogue, reflecting its conservative, clinically vetted design. Mycobacterium_pipeline detects substantially more loci outside the WHO catalogue (n=26), trading specificity for discovery potential. These trade-offs are not visible from any single self-reported number; they emerge only when the same benchmark is applied uniformly.

## 4 Community use and outlook

PathoBench launches with comprehensive coverage for *M. tuberculosis* as a worked example; for the other nine supported pathogens, the standard-dataset registry is complete and the framework is operational, but pipeline entries depend on community contribution. This is by design: a benchmark registry that pre-populates synthetic evaluations would inherit the very pathology — unverified self-report — it was built to address. We invite developers and laboratories worldwide to evaluate their tools against the listed datasets and submit results.

Three extensions are planned. First, real OAuth-based ORCID login will replace the current manual ORCID entry. Second, automated evidence checking will verify that submitted evidence URLs contain the claimed metric values. Third, annual reproducibility challenges, modelled on CAMI (Meyer et al., 2022), will encourage independent replication of high-impact submissions. Suggestions for additional pathogens and metrics may be filed as GitHub issues.

By making benchmarking a continuous, transparent, community-maintained activity rather than a periodic, fragmented one, we aim to give clinical and public-health bioinformaticians a reliable basis for tool selection — and tool developers a fair, visible venue for demonstrating their work’s contribution.

## Acknowledgements

We thank colleagues at the Bureau of Public Health Laboratories, Florida Department of Health, for testing the platform during development.

