## Supplementary Table S1 for "PathoBench: an open community-driven benchmark registry for pathogen bioinformatics tools"

**Supplementary Table S1. Pathogen-specific evaluation metrics in PathoBench**

*For each of the ten supported pathogens, PathoBench defines three to five evaluation dimensions that reflect that pathogen's analytical priorities. Submissions report all metrics as percentages (0–100); a composite score is computed as the unweighted mean across pathogen-specific metrics. Cross-pathogen comparison is intentionally not supported, as metric definitions differ between pathogens.*

| **Pathogen** | **Metric** | **Description** |
| --- | --- | --- |
| ***Mycobacterium tuberculosis*** | Drug coverage | Percentage of WHO-listed antimicrobials reported by the pipeline. |
|  | Gene coverage | Percentage of resistance genes in WHO catalogue covered by the pipeline. |
|  | Mutation confirmation | Percentage of pipeline-called mutations confirmable in WHO catalogue. |
| **HIV-1** | DRM concordance | Percentage of drug-resistance mutations matching Stanford HIVdb reference. |
|  | Drug-class coverage | Percentage of HIV drug classes (PI/NRTI/NNRTI/INI) reported. |
|  | Minor-variant sensitivity | Detection sensitivity for variants at 1–20% frequency. |
|  | Score concordance | Percentage agreement on Stanford-HIVdb interpretation scores. |
| **HBV** | Genotype accuracy | Percentage of correctly assigned HBV genotypes (A–J). |
|  | RT-resistance prediction | Percentage agreement on RT-inhibitor resistance interpretation. |
|  | HBsAg-escape detection | Percentage detection of clinically relevant S-gene escape mutations. |
|  | Recombinant handling | Percentage accuracy on recombinant / mixed-genotype samples. |
| **HCV** | Genotype accuracy | Percentage correct assignment of HCV genotypes 1–8. |
|  | Subtype resolution | Percentage correct subtype-level resolution within genotype. |
|  | RAS detection | Resistance-associated substitution detection vs reference panel. |
|  | Novel-variant handling | Performance on lineages absent from training reference. |
| **HAV** | Genotype accuracy | Percentage of correctly assigned HAV genotypes (I–VII). |
|  | VP1 coverage | Percentage coverage of the VP1/P2A diagnostic region. |
|  | Outbreak clustering | Concordance with reference phylogeny on outbreak clusters. |
| **SARS-CoV-2** | Lineage accuracy | Percentage correct Pango lineage assignments. |
|  | Clade accuracy | Percentage correct Nextstrain clade assignments. |
|  | Incomplete-genome robustness | Performance on partial / low-coverage sequences. |
|  | Version consistency | Stability of calls across tool versions. |
| **MPXV** | Clade accuracy | Percentage correct assignment to clades I, Ia, Ib, II, IIa, IIb. |
|  | Lineage resolution | Sub-clade lineage resolution. |
|  | APOBEC3 handling | Correct reconstruction in the presence of APOBEC3-driven mutations. |
|  | QC pass rate | Percentage of samples passing built-in quality control. |
| **Influenza** | Type accuracy | Correct identification of influenza A/B/C/D. |
|  | HA subtype | Correct HA subtype (e.g. H1, H3). |
|  | NA subtype | Correct NA subtype (e.g. N1, N2). |
|  | Antiviral resistance | Concordance with WHO-listed antiviral-resistance markers. |
|  | Mixed-infection detection | Sensitivity to mixed / co-infection samples. |
| **Ebola** | Species accuracy | Correct Ebolavirus species identification. |
|  | Sublineage resolution | Sub-clade resolution within an outbreak. |
|  | Phylogeographic concordance | Agreement with reference geographic phylogeny. |
|  | Real-time turnaround | End-to-end pipeline runtime (% of CDC reference baseline). |
| ***Plasmodium*** | Species accuracy | Correct identification of P. falciparum / vivax / ovale / malariae / knowlesi. |
|  | Geographic origin | Concordance with MalariaGEN geographic-origin assignments. |
|  | Drug-resistance markers | Detection of major resistance loci (Kelch13, Pfdhfr, Pfmdr1, etc.). |
|  | Co-infection detection | Sensitivity to mixed-species samples. |

*Metrics are dynamically defined per-pathogen in the platform source; updates may be proposed via the project's GitHub repository.*
